# A novel deep learning scheme for morphology-based classification of mycobacterial infection in unstained macrophages

**DOI:** 10.1101/611434

**Authors:** Xinzhuo Zhao, Yanqing Bao, Lin Wang, Wei Qian, Jianjun Sun

## Abstract

**Objective:** Mycobacterium tuberculosis (Mtb) is an airborne, contagious bacterial pathogen that causes widespread infections in humans. Using Mycobacterium marinum (Mm), a surrogate model organism for Mtb research, the present study develops a deep learning-based scheme that can classify the Mm-infected and uninfected macrophages in tissue culture solely based on morphological changes.

**Methods:** A novel weak-and semi-supervised learning method is developed to detect and extract the cells, firstly. Then, transfer learning and fine-tuning from the CNN is built to classify the infected and uninfected cells.

**Results:** The performance is evaluated by accuracy (ACC), sensitivity (SENS) and specificity (SPEC) with 10-fold cross-validation. It demonstrates that the scheme can classify the infected cells accurately and efficiently at the early infection stage. At 2 hour post infection (hpi), we achieve the ACC of 0.923 ± 0.005, SENS of 0.938 ± 0.020, and SPEC of 0.905 ± 0.019, indicating that the scheme has detected significant morphological differences between the infected and uninfected macrophages, although these differences are hardly visible to naked eyes. Interestingly, the ACC at 12 and 24 hpi are 0.749 ± 0.010 and 0.824 ± 0.009, respectively, suggesting that the infection-induced morphological changes are dynamic throughout the infection. Finally, deconvolution with guided propagation maps the key morphological features contributing to the classification.

**Significance:** This proof-of-concept study provides a novel venue to investigate bacterial pathogenesis in a macroscopic level and has a great promise in diagnosis of bacterial infections.

## I. Introduction

MYCOBACTERIUM is a genus of Actinobacteria [1], which includes pathogens known to cause serious diseases in humans, such as Mycobacterium tuberculosis (Mtb) that causes global tuberculosis epidemic and accounts for more than 15 millions of deaths since 2000 [2]. Early and accurate detection of Mtb infection is an urgent need. Traditional diagnosis of TB relies on recovery of live Mtb organisms from patients’ sputum samples by microbiological culture, but this method is time consuming and has a low detection rate [3]. Recently, various molecular and immunological methods have been developed for TB diagnosis, including sputum smear microscopy [4][5], tuberculin skin test [6], interferon-γ release assays [7], and Xpert MTB/RIF [8], etc. Although each of these methods has unique advantages and has been used in some clinical settings, none of them can directly detect live Mtb organisms from patients. Thus the traditional microbiological culture of live Mtb from patients remains as the well-accepted gold standard for TB diagnosis.

Image analysis has a great potential in disease diagnosis. Large amounts of advanced quantitative features from images are extracted and analyzed, and the image data are processed in a mineable form to build descriptive and predictive models correlating image features to specific diseases even with gene-protein signatures [9]. For instances, medical image analysis has been able to distinguish benignant lung nodules from the malignant ones by the CT scan images [10][11]. Breast tumors can be detected and analyzed automatically based on the radiological images [12]–[14]. The brain partitions have been segmented for further exploration [15][16]. Image analysis is also extensively involved in basic biomedical sciences and biology. For instances, the cell organs are segmented [17] and cellular classification tasks are also manipulated by image analysis [18].

There are two main challenges for the image analysis: feature extraction and the choice of the classifier. For the feature extraction, deep learning, especially the convolutional neural network (CNN), has made a dramatic progress [19]. Compared with traditional hand-crafted features, CNN extracts ample features automatically. In natural images classification task, CNN has achieved superhuman performance [20]–[22]. Generally, the CNN model can be generated by training from scratch or from the transfer learning method. Transfer learning means to utilize the knowledge learned from one task to another task, as long as the two tasks share the same features [23]. Compared with training the CNN model from scratch, transfer learning involves less manual modifications and less input images. It also helps to enhance the classification performance and reduce the training time. For the classifier, there are three kinds of machine learning methods: unsupervised, semi-supervised, and supervised learning. Unsupervised learning trains the predictive model with unlabeled data. For example, images features can be related to gene-mutation by unsupervised learning without any prior knowledge [24]. Supervised learning generally obtains more accurate results, owning to its abundant labeled data training process. Semi-supervised learning falls between, which requests only a small amount of labeled data with a large amount of unlabeled data. Thus, it combines the advantages of unsupervised and supervised learning methods.

Considering the significant success of image analysis in medical and biological sciences, we are inspired to introduce CNN to the classification task of the microscopic world. In this study, we systematically design a scheme to distinguish mycobacteria-infected cells from uninfected cells based on cellular morphological changes that are indistinguishable to naked eyes at the early stage of infection. First, a weak-and semi-supervised method is developed to detect cells from the images. It combines the gradient-based circle Hough transform (GBCHT) algorithm [25] with the supervised deep learning. Second, the cells are classified by transfer learning and fine-tuning. Third, image reconstruction by deconvolution is applied to detect the key morphological features contributing to the classification results.

## II. Materials and Methods

### A Data Acquisition

Mycobacterium marinum (Mm) is a pathogen that causes tuberculosis-like diseases in fish and sometimes causes mild skin infections in human. Since Mm shares similarities with Mtb, in genomes, virulence factors and modes of infection in macrophages, it is widely accepted as a safer surrogate model for Mtb in laboratory research. Here, we use a Mm strain harboring a pMSP12:mCherry plasmid (Addgene, USA), which expresses a mCherry protein so that the bacteria emit cherry fluorescence [26]. Thus, the cherry fluorescence serves as an infection marker to train CNN and also to validate the classification results.

At the early stage of Mtb infection, alveolar macrophage is the first phagocyte that recognizes and engulfs Mtb. The murine macrophage cell line RAW264.7 is one of the most commonly used cell line for the studies of intracellular infection by Mtb and Mm. Thus, this study uses RAW264.7 cells as the infected host model.

RAW264.7 cells are plated in 8-well chambered culture slides (Corning Falcon, USA) at 1×10^5^ cells/well and cultured in Dulbecco’s modified eagle medium (DMEM, Hyclone, USA) containing 10% fetal bovine serum (FBS, Corning, USA) at 37°C, 5% CO2 for 24 hr. Cells are washed three times with pre-warmed PBS to remove residual antibiotics. Then, the Mm bacteria expressing mCherry are added into the wells to reach 2 multiplicity of infection (MOI), followed by centrifugation at 250 × g at room temperature (RT) for 5 min. After 30 min incubation at 30°C, the cells are washed again with pre-warmed PBS to remove unbound bacteria. Then DMEM with 100µg/ml Amikacin is added to kill the extracellular bacteria for 1 hr at 30 °C. The infected cells are cultured in DMEM containing 1% FBS, 50µg/ml Amikacin at 30°C, 5% CO_2_ until image acquisition.

At 2, 12 and 24 hours post infection (hpi), the cells are washed with PBS and fixed with 4% paraformaldehyde for 10 min at RT. Once the fixed slides are dried in air, mount media (Enzo Life Sciences, USA) is added to prevent fluorescence quench. Finally, the slides are covered with cover slips. The slides are imaged with a FLoid Cell Imaging Station (Life technologies, USA) at 460 X magnification with the white light channel to image the cells and the red fluorescence channel to image the cherry color bacteria. The images were exported as 16-bit tagged image file format (TIFF) with a final resolution of 96 digital pixels per inch (dpi).

### B Cell Detection

All of the image processing and analysis are performed using MATLAB language and Caffe [27] framework in a workstation with a GTX1060 GPU.

A novel weak-and semi-supervised detection (WSSD) algorithm is developed for cell detection. This method combines the gradient-based circular Hough transform (GBCHT) [25] with the convolutional neural network. GBCHT is a feature-based learning method and doesn’t need manually labeled input data. CNN is a supervised learning algorithm. Our novel algorithm uses a small amount of manually labeled data along with a large amount of unlabeled data, which is therefore named as weak-and semi-supervised detection (WSSD) algorithm.

Hough transform [28] was put forward to detect specific line patterns. When detecting circles, the points in the images are transformed to the parameter space, seen as (1):

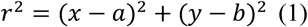

leading the point represented by three parameters: *r, a* and *b*. Where *r* is the radius of each detected circle, and *a* and *b* present the location of the center of the circle. To improve the detection accuracy, gradient is utilized in [25][29]. First, the gradient to the center of the whole image is calculated by (2):

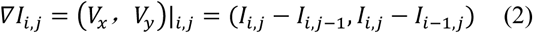

where (*i, j*) indices the pixel position. *I*_*i,j*_ is the intensity of the pixel. ∇*I*_.,0_ is the gradient vector of the pixel at the position *(i, j)*. Then, an accumulation step is introduced to find the highest response of the gradient, which would be the center of the circle. For the determination of the radii [30], if the point satisfies (3), where Δ*r* is the interval between adjacent *r* values.

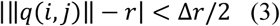

Besides cell detection, small amounts of cells are labeled as the positive or negative cells in instance-level (i.e., bounding boxes), as shown in Fig. 1. Instead of classifying the cells in the image-level labels, it’s preferred to distinguish which single cell is infected. Because in the images of the infected group, not all of the cells are infected. Then, these labeled cells are used to train a classifier to discriminate the true infected cells from the uninfected cells. Transfer learning and fine-tuning from the CNN are utilized, which will be introduced in next subsection. After the classifier is well trained, the rest of the cells, which are the unlabeled ones, are classified by the prediction model.

**Fig. 1.**
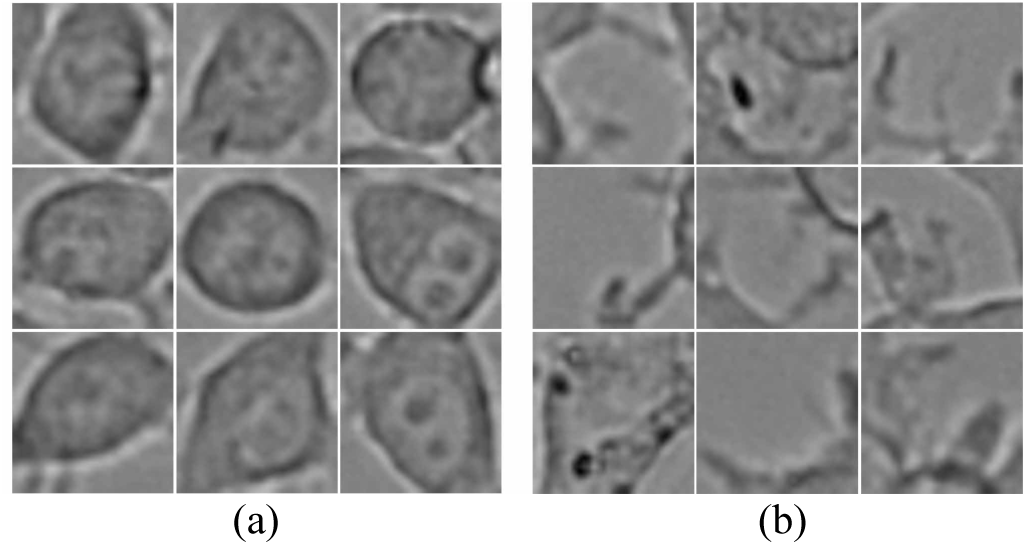
Manually labeled cells of the GBCHT detection. The cells are initially detected from the whole images by the GBCHT algorithm. Considering there being miss detected or error detected cells, the detected cells are labeled as the positive cells, shown in (a), and the negative cells, shown in (b). The labeled cells are then utilized for further supervised learning process.

### C Transfer Learning and Fine-tuning for Classification

This is a binary classification task. There are two experimental groups: the control group, in which all the cells are not infected; the infection group, which contains infected cells and uninfected cells since the rate of infection is less than 100%. The cells in control group are extracted directly by WSSD as the labeled uninfected cells. For the infection group, only the cells with cherry color bacteria either within or on the circles are considered as infected cells and extracted from the images, as seen in Fig. 2.

**Fig. 2.**
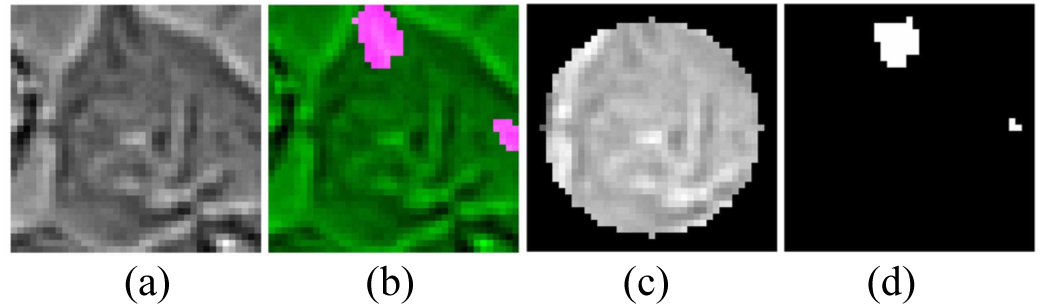
Extraction of the Mm-infected cells. Since the rate of infection is less than 100%, extra filter is applied to remove the cells without mCherry fluorescence (regarded as uninfected cells). (a) an infected cell in the view of white light; (b) the merge of mCherry fluorescence from bacteria with the white light images; (c) the mask for the cell; (d) the overlapping region between the cell and the bacteria.

#### 1) Convolutional neural network architecture

ResNet [31], a contemporary CNN architecture, is one of the best deep learning classifier with 152 layers. It is designed for 224×224 pixel images. In view of its deep structure, the memory of GPU can’t meet the computational cost. Thus, a cropped version with ResNet-50 [32] is utilized in this experiment. Seen in Table I, it’s a stack of 5 groups of convolutional layers. Each group consists of several residual blocks. It adds on a “delta” or a slight change to the original input, called “residual”.

**TABLE I.**
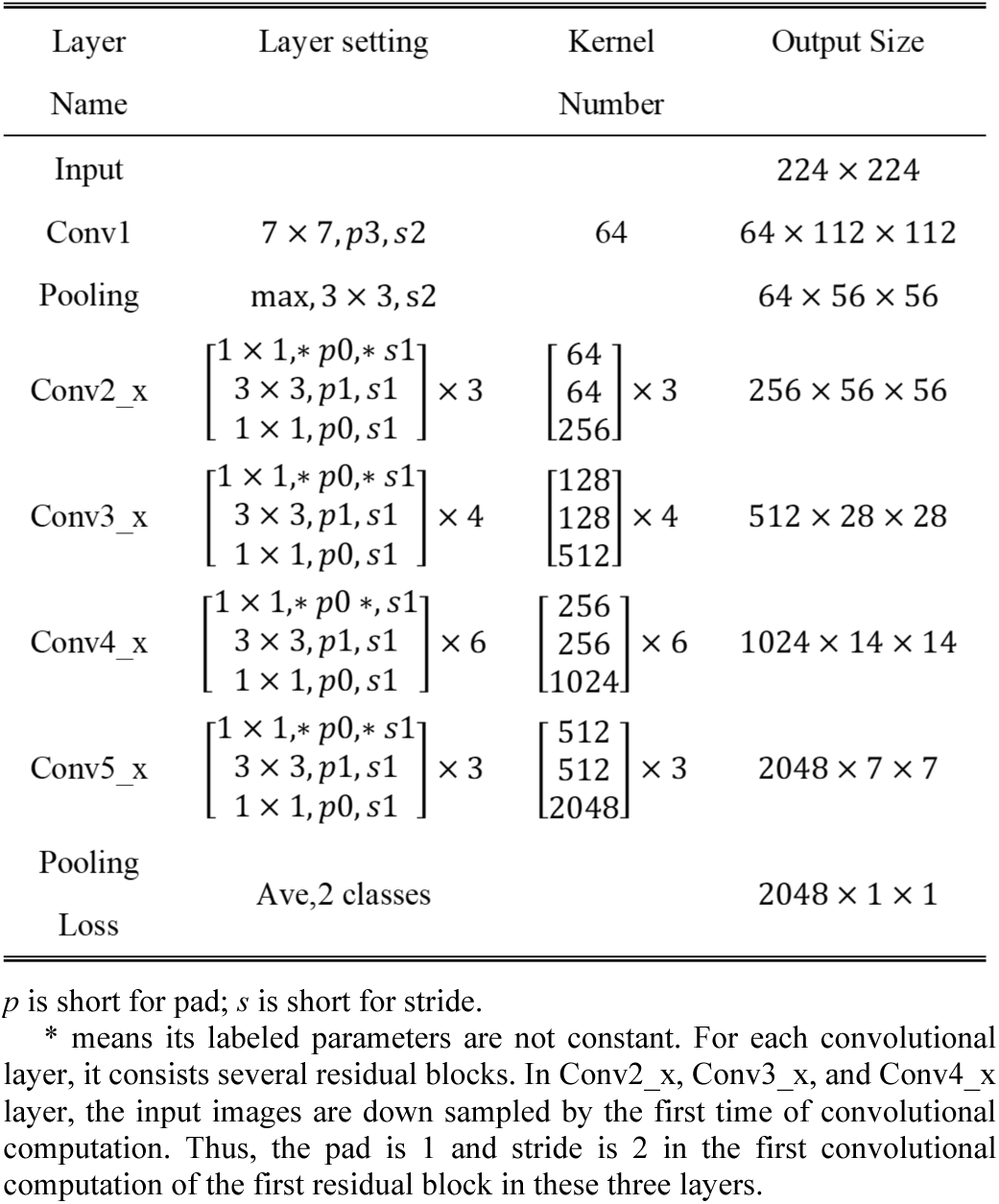
The Structure of the Transfer Learning and Fine-Tuning from the Reset-50

#### 2) Transfer learning and fine-tuning from CNN

There are three methods to implement transfer learning and fine-tuning. (1) Only transfer learning without fine-tuning. The trained deep learning model is directly used for prediction without adjustment by the new task images. This method is underperforming. (2) Transfer learning with fine-tuning of the last classification layer. For example, the trained model is based on the ImageNet that classifies the 1000 classes. The new task just needs to distinguish between 10 classes. This method needs to initialize the parameters of the last classification layer and to change the output class into 10. The former convolutional layers do not participate the backpropagation, which will not be fine-tuned. Compared with the other two methods, this one performs in between. (3) Transfer learning with fine-tuning of all the layers, including the convolutional layers. It means all the weights inherited from the trained deep learning model are fine-tuned by the new task datasets and all layers participate the backward computation. This one outperforms the other two [33]. In our experiment, the third method, which performs the best, is utilized.

At the beginning, the pre-trained network is trained by the natural images with random initialized weights. Then, the pre-trained model is transferred to Mm infection classification task. The weights and bias of the last fully connected layers are initialized randomly. The neural units are reduced to two. None of any layers in the entire network have frozen the weights and bias. All of them are fine-tuned by backward computation. The learning rates are changed to 0.0005, which is far smaller than their original one.

### D Deconvolution

Deconvolution is a method for visualizing the features learned by the hidden units of CNNs. It’s also named as transposed convolution or backward stride convolution[34]. In deep learning, it is an inverse flow of its original training process. Given a high level feature map, by the backward computation, the reconstructed image will show the strongly activated part of the input images. By deconvolution, the hidden layers of deep neural network can be observed by eyes. Zeiler introduced a novel visualization technique that gave insights into the function of intermediate feature layers and analyzed the strongest activation of the input images by deconvolutional network. The observation of the deconvolutional results is improved by the guided backpropagation [35].

In this experiment, guide backpropagation is performed to reconstruct image. In the training process, the input images are down sampled by several convolutional layers and pooling layers. Therefore, when reconstructing the images, two up sampling ways are proceeded. By adjusting the stride steps or padding numbers, the reconstructed images can be up sampled. The details are well described in [36]. Beside backward stride convolution, up-pooling is another key step to affect the final reconstruction result. There are three main methods for reconstruction: back propagation, deconvnet and guide backpropagation. When doing the ReLU of forward propagation, the position of eliminated pixels (less than zero) will be recorded. In back propagation, the same position of the pixels will still be eliminated from the backpropagation, regardless of its value being less than zero or not. In deconvnet [37], the position of eliminated pixels will be reevaluated by the actual value (whether is less than zero). Guided backpropagation combines these two methods, which removes both of the values eliminated by the last two ones. The deconvolution reconstruction is based on the DeepVis Toolbox [37].

## III Results

### A. The Comparison between WSSD and GBCHT

The cell detection results of GBCHT are shown in Fig. 3. The original cell image is exhibited in Fig. 3a and the accumulation array of each pixel’s gradient is shown in Fig. 3b. By searching the local maxima, the center of the circle can be detected. Although, not all the cells are regular circular, both circular and irregular shapes can still be observed in the experiment, as seen in Fig. 3c. The red cross in the middle of the image is supposed to be the circle center. Besides the irregular shapes, the algorithm is also robust to the overlapped cells, as seen in Fig. 3d. They are also correctly detected in Fig. 3f. To evaluate the detection performance, quantitative assessment is implement. Some cells are correctly detected (Fig. 3f) and named as true positive (TP). The yellow squared cell shown in the Fig. 3c is a cell that is missed in detection, named as false negative (FN). Besides those, some non-cell regions were detected as cells by error, named as false positive (FP), as seen in Fig. 3g.

**Fig. 3.**
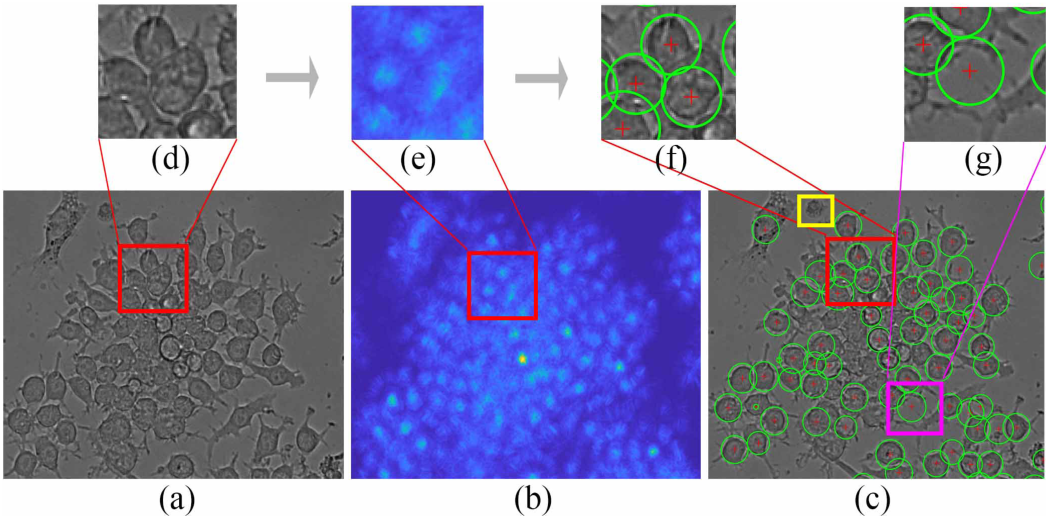
Cell detection results with gradient-based circular Hough transform. (a) The original image taken by white light channel; (b) The accumulation array of each pixel’s gradient. The high response points are the detected centers of the circles; (c) The detection results with each green circle indicating a cell. The yellow square shows a cell that is missed in detection. To observe the details, parts of the images are enlarged and shown as subgraphs. (d) – (e) The example of the overlapped cells. (g) The error detected example.

To further improve the detection performance, WSSD is developed. We use the same images to evaluate the performance of WSSD, for the comparison between GBCHT. The statistical results are shown in Table II. It’s obvious that there are significant improvements of the new method in accuracy and precision with *p* < 0.05. According to the results, this new method reduces the false positive. Granted that, the WSSD’s sensitivity is not as high as GBCHT’s, meaning more real cells are misidentified as the background, but it can still ensure that the 99.4% remaining cells are the true positive ones.

**TABLE II.**
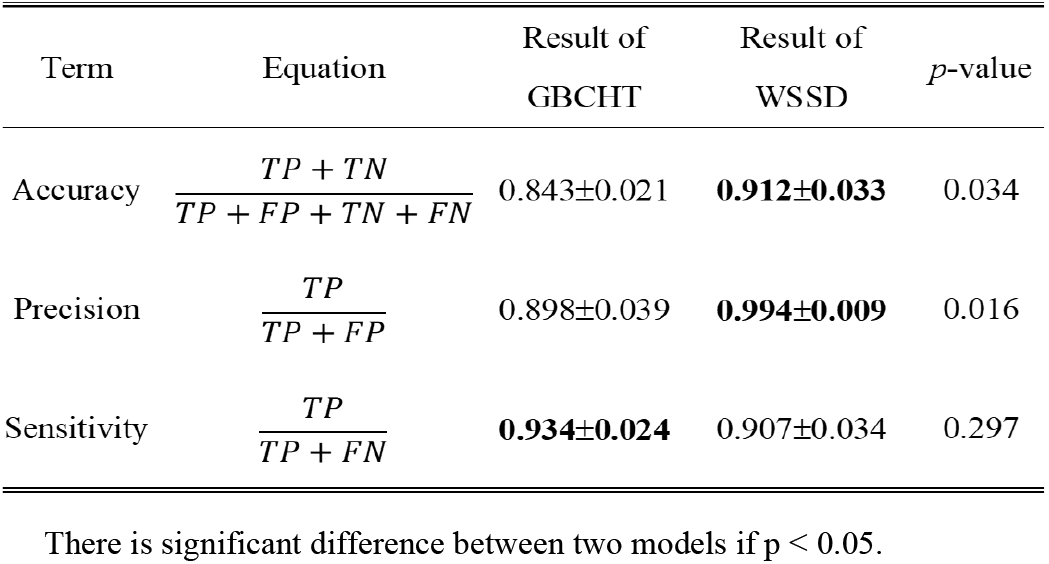
The Summary of the Detect Results

### B. Evaluation of the Cell Classification

Transfer learning and fine-tuning from ResNet are used to distinguish the uninfected cell from the Mm-infected cells at 2, 12, and 24 hpi. Fig. 4 presents the training loss and the validation accuracy at 2 hpi. The training loss is reduced with the iteration. And the validation accuracy increases quickly with the iteration, which reaches to 90% average accuracy. To estimate the “health” of the network, feature maps are observed. Three of them are shown in Fig. 5, which are conv1, conv2 (layer_64_2_conv2), and conv4 (layer_256_2_conv2), respectively. These features show the complex invariances learned in the convolutional layers. The lower layer, seen as the Fig. 5a, shows an organized structure pattern, which are understandable to human. These basic textures indicate the network to be qualified to detect the structure edges of the input images. The higher layer, seen as Fig. 5b and 5c, shows some incomprehensible features. Their uncorrelated patterns ensure the variation of different channels, which also guarantees a “healthy network”

**Fig. 4.**
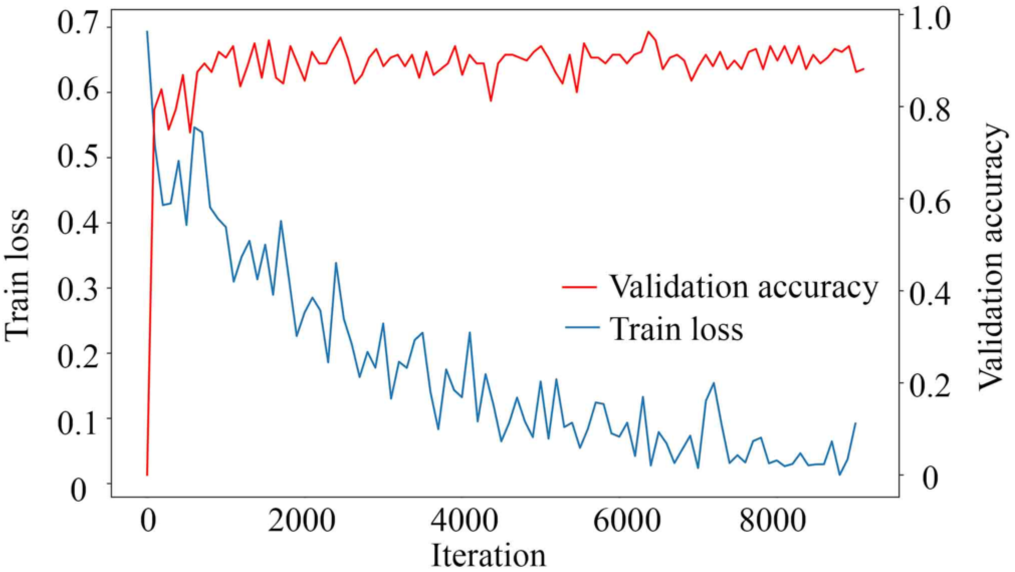
The train loss and validation accuracy of the transfer learning. The blue line presents the training loss changing with the iteration. The red line shows the validation accuracy.

**Fig. 5.**
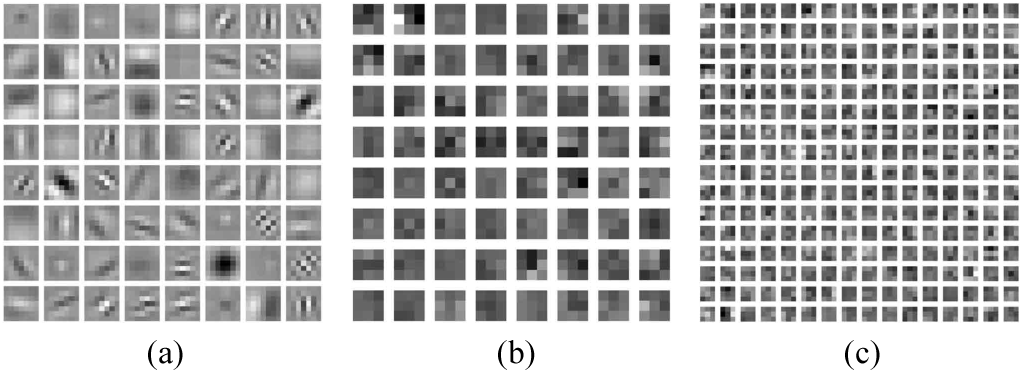
Feature examples of the three layers from the trained CNN for classification. (a) is the features from the conv1 layer; (b) and (c) are features of conv2 (layer_64_2_conv2) and conv4 (layer_256_2_conv2) layer. Structured and heterogeneous patterns indicate a well-trained CNN model.

The performance of classification is evaluated using the 10-fold cross-validation. The results are summarized in Table III It summarizes the total number of each group of cells and the classification results. Seen as the second and the third column, the numbers of two classes of each time are almost balanced. This balance leads no additional bias to the results. The classification results are also exhibited in an intuitional way in Fig. 6. There are distinct differences between the uninfected cells and Mm-infected cells at 2 hpi, which achieves the ACC of 0.938 ± 0.020. However, at 12 hpi, the cells of two classes are difficult to distinguish, achieving the ACC of 0.749±0.010. It indicates less difference in morphologies of two classes of cells. At 24 hpi, the ACC turns out to be 0.824±0.009. SENS and SPEC have the same trends as the ACC.

**TABLE III.**
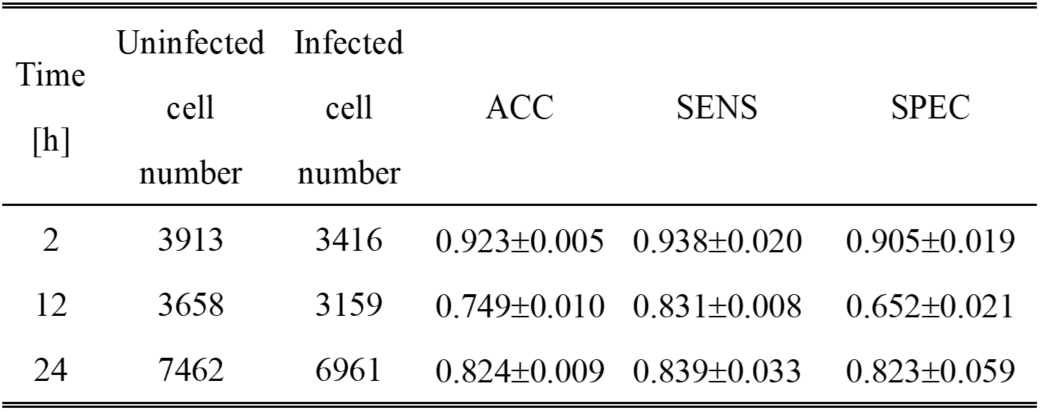
Cell Numbers and the Classification Results

**Fig. 6.**
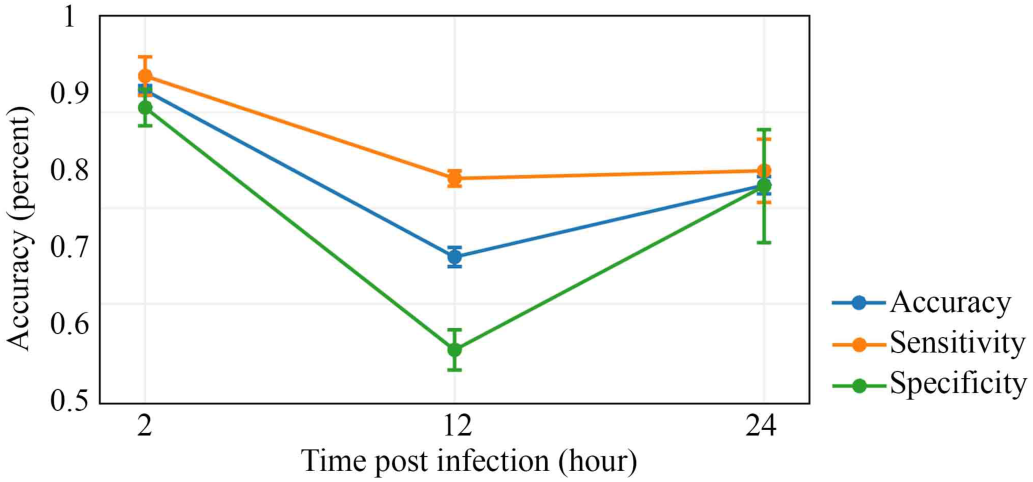
Classification results of uninfected cells and Mm-infected cells. The cells from infected group or uninfected control group were classified, and Accuracy (blue), Sensitivity (orange) and Specificity (green) were calculated and plotted against the hours of post-infection.

### C. Guided backpropagation for deconvolution

Guided backpropagation is utilized to analyze the influence of each pixel to the classification results. The deconvolutional results of the uninfected control cells (Fig. 7a) and infected cells (Fig. 7b) at 2 hpi are presented. Three guided backpropagation examples from each group are selected because their channels are highly activated, which means they contribute significantly to the decision of the classification. The high active signals in both groups focus on the cell edges, suggesting that cell edges are significant features in the classification of two groups of cells. It is also observed that there are more dots inside the infected cells, such as the bottom right images (Fig. 7b). More interestingly, the high activated pixels have little overlap with the cherry color bacteria, indicating that the CNN does not classify the cells by detecting the bacteria.

**Fig. 7.**
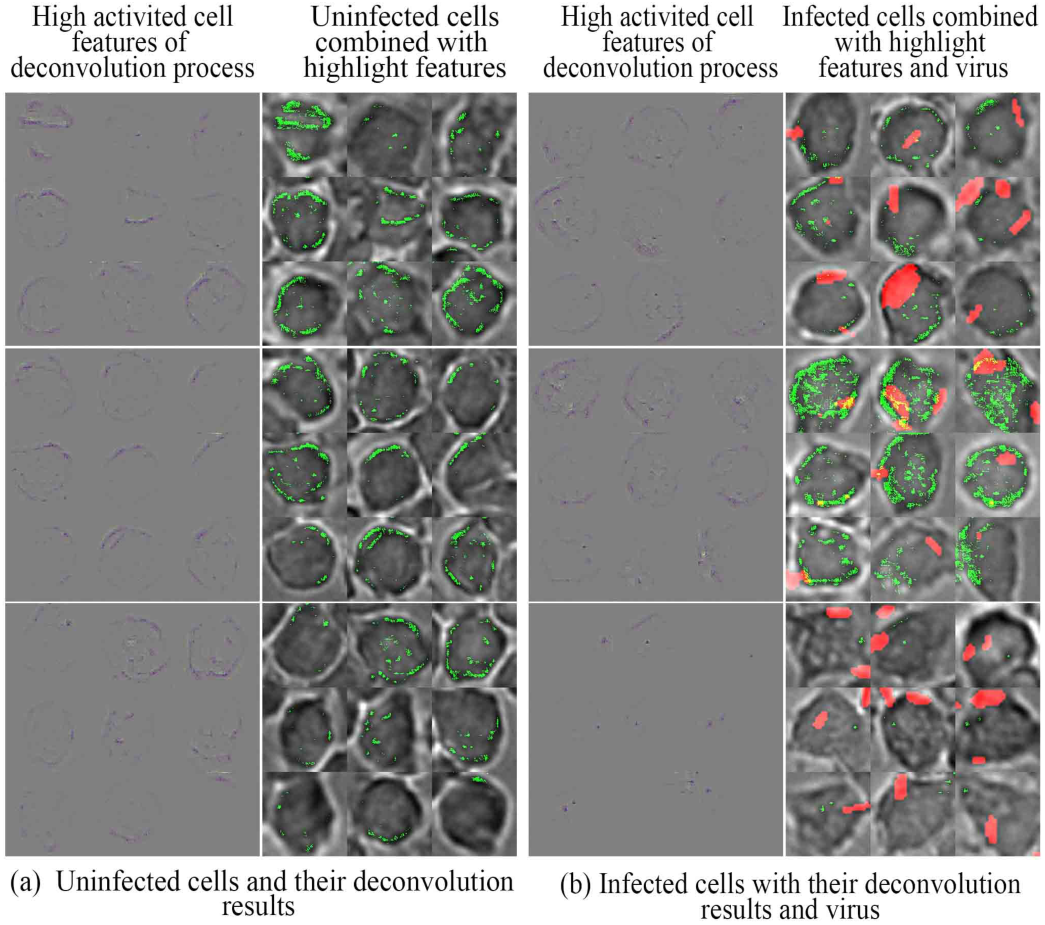
Guided backpropagation results of the cells from the uninfected control group and Mm-infected group. Each group contains three samples shown in rows. Each sample is presented by two versions: deconvolutional results (left column) and the original cell images merged with the deconvolutional results (right column). The deconvolution signals are presented in green. The control group is shown in (a). The Mm-infected group is shown in (b), in which red regions indicate the bacteria.

## IV. Discussion

This study is the first time that deep learning-based image analysis is used to detect Mm infection in cultured macrophages. This method detects significant morphological differences between the infected and the uninfected cells as early as 2 hpi, even though these changes are indistinguishable to naked eyes. Interestingly, the detection accuracy varies at different times of infection. It is high at 2 and 24 hpi and become significantly lower at 12 hpi. This result indicates that the host-pathogen interaction is a dynamic process and the infection-induced morphological changes vary accordingly in different infection stages. In fact, it is correlated surprisingly well with the widely-accepted experimental data that at the early stage of infection, macrophages make strong immune responses to mycobacteria, including engulfing mycobacteria into the phagosomes and activating oxidative stress within the phago-lysosomes [38][39], while at the same time, mycobacteria exert various mechanisms to counteract and suppress the host immune responses, which should be accompanied with significant morphological changes until a balance between host and mycobacteria or a latency is established at the middle stage of the infection [40]–[42]. At the later stage of infection, mycobacteria may break the latency state and transition to active infection, which is evidenced by the observation that Mm escapes the phagosome into the cytosol, where Mm replicates, recruits actin tails for cell-to-cell spreading, and eventually causes cell death [43]–[45]. The CNN-detected morphological changes, which are invisible to naked eyes, provide a novel venue to investigate into the mechanism of bacterial pathogenesis.

The newly developed method has three advantages in cell detection: robustness, effectiveness and preciseness. The original GBCHT is widely used in various areas, such as to identify the eyes in facial images [46] or to detect multiple object instances [47]. The developed WSSD doesn’t need parameter setting, making it accessible to various tasks. Supervised learning method generally outperforms unsupervised method with the cost of large amount of manual labeled data. For instance, U-Net is a state-of-the-art architecture for detection and segmentation of cells in light microscopy images [48]. Though it achieves good results, it is less favored because of labor consuming. However, in our method, only small amounts of labeled data are needed. The semi-supervised method is efficient because it demands less labeled data. It is also more accurate than the unsupervised method. Currently, this method is only sensitive to quasi-circle objects. Thus, some cells are missed due to irregular shapes, such as extensional pseudopodia. However, it doesn’t affect the results as long as enough true positive cells are extracted for analysis, which meet our goal. Taking all these factors into account, the WSSD is well developed in cell detection.

While some cell features are well extracted and analyzed by hand-crafted methods [49]–[51], deep learning has proven to have supreme performance in medical image diagnosis [52]. Xu et al. extracted features of the sickle cell and classified with CNN [53]. Chen et al. also classified the cells by heuristic genetic algorithm using the extracted features [54]. Different from these two works, our raw data are fed directly to the CNN is more sufficient than those designed by human hands. What’s more, among the contemporary CNN structures, ResNet outperforms most of the other ones, such as AlexNet [20], VGGNet [21], and GoogLeNet [55]. Therefore, we build our architecture based on the ResNet-50. Proven by our experiment results, the classification accuracy at 2 hpi is as high as 0.923±0.005.

The deep learning method accurately detects the Mm-induced morphological changes that are indistinguishable to human eyes. It is important to understand where those changes are located and what cellular structures are affected, which will provide clues for future studies to reveal the in-depth mechanism of host-pathogen interaction.

## V. Conclusion

In summary, this study has successfully developed a deep learning-based scheme to classify mycobacteria-infected cells from the uninfected cells. The cells are detected and extracted with a weak-and semi-supervised detection algorithm. Excellent classification accuracy is achieved by the transfer learning and fine-tuning of the CNN. Deconvolution reconstructs the CNN-detected morphological features, which will provide a novel venue in studies of microbial pathogenesis. This scheme opens a new route to not only detect bacterial infection but also to other biological tasks.

## Notes

Research reported in this publication was supported by the National Institute Of General Medical Sciences of the National Institutes of Health under Award Number SC1GM095475 (to J. Sun), National Center for Research Resources (5G12RR008124); and National Institute on Minority Health and Health Disparities (G12MD007592).. The content is solely the responsibility of the authors and does not necessarily represent the official views of the National Institutes of Health

